# Postdiction: when temporal regularity drives space perception through pre-stimulus alpha oscillations

**DOI:** 10.1101/2021.01.24.427978

**Authors:** Laetitia Grabot, Christoph Kayser, Virginie van Wassenhove

## Abstract

During postdiction, the last stimulus of a sequence changes the perception of the preceding ones: in the *rabbit* illusion, a three-flash series presented regularly in time, but not in space, is – illusory - perceived as spatially regular. Such a reorganization of (spatial) perception could be driven by internal priors, e.g. favoring slow motion for the *rabbit* illusion. Although postdiction is a ubiquitous phenomenon, its neural underpinnings remain poorly understood. Here, we focused on the role of priors during postdiction and hypothesized that these could be reflected by alpha oscillations (8-12Hz), previously observed to correlate with idiosyncratic biases. We presented human participants with ambiguous visual stimuli that elicited the *rabbit* illusion on about half the trials, allowing us to contrast MEG-EEG brain responses to the same physical events causing distinct percepts. Given that a strong prior will increase the overall probability of perceiving the illusion, we used the percentage of perceived illusion as a proxy for an individual’s prior. We found that high fronto-parietal alpha power was associated with perceiving the sequence according to individual biases: participants with high susceptibility to the illusion would report the illusion, while participants with low susceptibility would report the veridical sequence. Additionally, we found that pre-stimulus alpha phase in occipital and frontal areas dissociated illusory from non-illusory trials. These results point to a dissociated relation of the power and timing of alpha band activity to illusory perception, with power reflecting prior expectations and phase influencing behavioral performance, potentially due to the modulation of sensory uncertainty.

**Significance Statement:** Late events may sometimes influence how earlier events are perceived, as if the arrow of time was reversed in the brain. This surprising phenomenon, called postdiction, is observed in the rabbit illusion, and highlights a predominant mechanism for perceptual processes. Perception builds up from the combination of prior expectations with incoming sensory evidence, which takes time. We showed that pre-stimulus neural activity, and more specifically alpha oscillations (8-12Hz), play a double role in postdiction. Fronto-parietal alpha power reflects individual prior expectation, while occipital and frontal alpha phase predicts illusory perception. Postdiction might actually be a means of compensating for the neural delays inherent in perceptual processes, so that the arrow of perceptual time matches the arrow of physical time.

## Introduction

Our phenomenological awareness of the world arises from the confrontation between sensory inputs that unfold over time, and the predictions derived from internal models the brain uses to infer causality between sensory events. Here, we investigated how the content of consciousness is updated, or more specifically, how late events can modify the perception of earlier events. In the *rabbit* illusion, stimuli in a sequence can causally affect the perceived localization of earlier events (Geldard & Sherrick, 1972): When three flashes are presented at a regular rate in the visual periphery, the first two at the same location and the last at a different one, participants often mislocalize the second flash to be in between the first and the last flash. This illusion, one of the rare true temporal illusions (Dennett & Kinsbourne, 1992), provides a means to study postdiction (Eagleman & Sejnowski, 2000), which may constitute an ubiquitous mechanism shaping perception (Shimojo, 2014).

To date, only one fMRI study and a preliminary EEG study have investigated the neural correlates of postdiction (Blankenburg, Ruff, Deichmann, Rees, & Driver, 2006; Stogbauer, van Wassenhove, & Shimojo, 2007), leaving clear gaps in our understanding of the underlying neural mechanisms. Computational and behavioral studies proposed that the reorganization of temporally related information, occurring during the *rabbit* illusion, could be driven by internal priors favoring slow-speed visual motion (Jones & Huang, 1982; Goldreich & Tong, 2013; Khoei, Masson, & Perrinet, 2017). The aim of the present study was to investigate the neural processes reflecting the impact of such priors during postdiction. We addressed this by recording simultaneous magneto- and electro-encephalography (MEEG) brain activity in human participants, allowing for both a fine temporal and spatial localization of relevant sources (Henson, Mouchlianitis, & Friston, 2009).

Previous work suggested that potential correlates of the impact of priors on behavior are visible in spontaneous brain activity, such as alpha band oscillations (Mayer, Schwiedrzik, Wibral, Singer, & Melloni, 2016; Samaha, Boutonnet, Postle, & Lupyan, 2018). In fact, many studies have shown that alpha oscillations correlate with the perception of upcoming stimuli (Ergenoglu, et al., 2004; Hanslmayr, et al., 2007; van Dijk, Schoffelen, Oostenveld, & Jensen, 2008; Mathewson, Gratton, Fabiani, Beck, & Ro, 2009). Increased alpha power was notably associated with a stronger influence of idiosyncratic biases of individual observers (Grabot & Kayser, 2019; Grabot, Kösem, Azizi, & Wassenhove, 2017). Since these biases may come from individual prior expectation (de Lange, Heilbron, & Kok, 2018), we hypothesized that pre-stimulus alpha power would reflect the impact of the intrinsic prior involved in postdiction. To measure these intrinsic priors, we used inter-individual differences in the tendency to perceive the illusion: for a given sequence of stimuli, some participants are more prone to perceive the illusion, whereas others are more likely to perceive the veridical sequence. Those differences may be due to inter-individual differences in the strength of internal priors. We hypothesized that pre-stimulus alpha power predicts an individual’s tendency to perceive the illusion. For comparison, we also tested the alternative hypothesis proposing that alpha relates to accuracy (Ergenoglu, et al., 2004; Mathewson, Gratton, Fabiani, Beck, & Ro, 2009), by directly contrasting pre-stimulus alpha power between illusory and non-illusory trials.

To complement the above analysis of oscillatory power, we also investigated the role of pre-stimulus phase in postdiction, since low-frequency phase were reported to influence the perception of upcoming stimulus (Busch, Dubois, & VanRullen, 2009; Mathewson, Gratton, Fabiani, Beck, & Ro, 2009) and illusory perceptions (Dugué, Marque, & VanRullen, 2011; Samaha & Postle, 2015; Chakravarthi & VanRullen, 2012; Keil, Schnitzler, Dijk, Weisz, & Lange, 2014). Given the literature (VanRullen, 2016) and the temporal delay of the visual sequence (<200ms) eliciting postdiction (Geldard, 1982), we focused on the alpha (8-12 Hz) and theta (4-7 Hz) bands. We hypothesized that low-frequency phases can dissociate illusory from non-illusory percepts.

We found that pre-stimulus parietal and frontal alpha activity fluctuations correlate with the individuals’ tendency to perceive the illusion, suggesting that alpha reflects the strength of the influence of an internal prior. Alpha phase in occipital and frontal areas was also predictive of whether a sequence would be perceived as illusory or veridical.

## Materials and Methods

### Participants

19 right-handed participants (8 males; age = 28 years, SD = 6) took part in the study. All had normal or corrected-to-normal vision, were under no medical treatment, no known neurological history, and were naive as to the purpose of the study. Each participant provided a written informed consent in accordance with the Declaration of Helsinki (2013) and the Ethics Committee on Human Research at NeuroSpin (Gif-sur-Yvette, France). Six subjects were excluded a priori from the analysis due to the failure of the digitization procedure (1 participant), data which were too noisy (1 participant), three subjects did not reach the 60% performance criterion in the control task (for one or two conditions) and one subject did not have enough trials in one condition. Hence, 13 participants (6 males, age = 29 years, SD = 7) were considered for MEG analyses.

### Experimental design

The visual stimuli were M-scaled 2D Gaussian blobs presented in a sequence of three flashes. The visual sequence was located on the right horizontal meridian with regularly spaced locations at 15.1°, 18.5° and 21.8° of visual angle. To account for visual acuity differences according to the stimulus position to fovea (Duncan, 2003), the diameter of the flashes was 2.5°, 3° and 3.5°, respectively. Each flash was presented for 50 ms. In the control condition, the second flash was presented at an intermediate position between the first and the last flash in the sequence, constituting a regular spatiotemporal sequence. In the test condition, the second flash was presented at the same position as the first flash in the sequence (Fig. 1A).

**Fig. 1.**
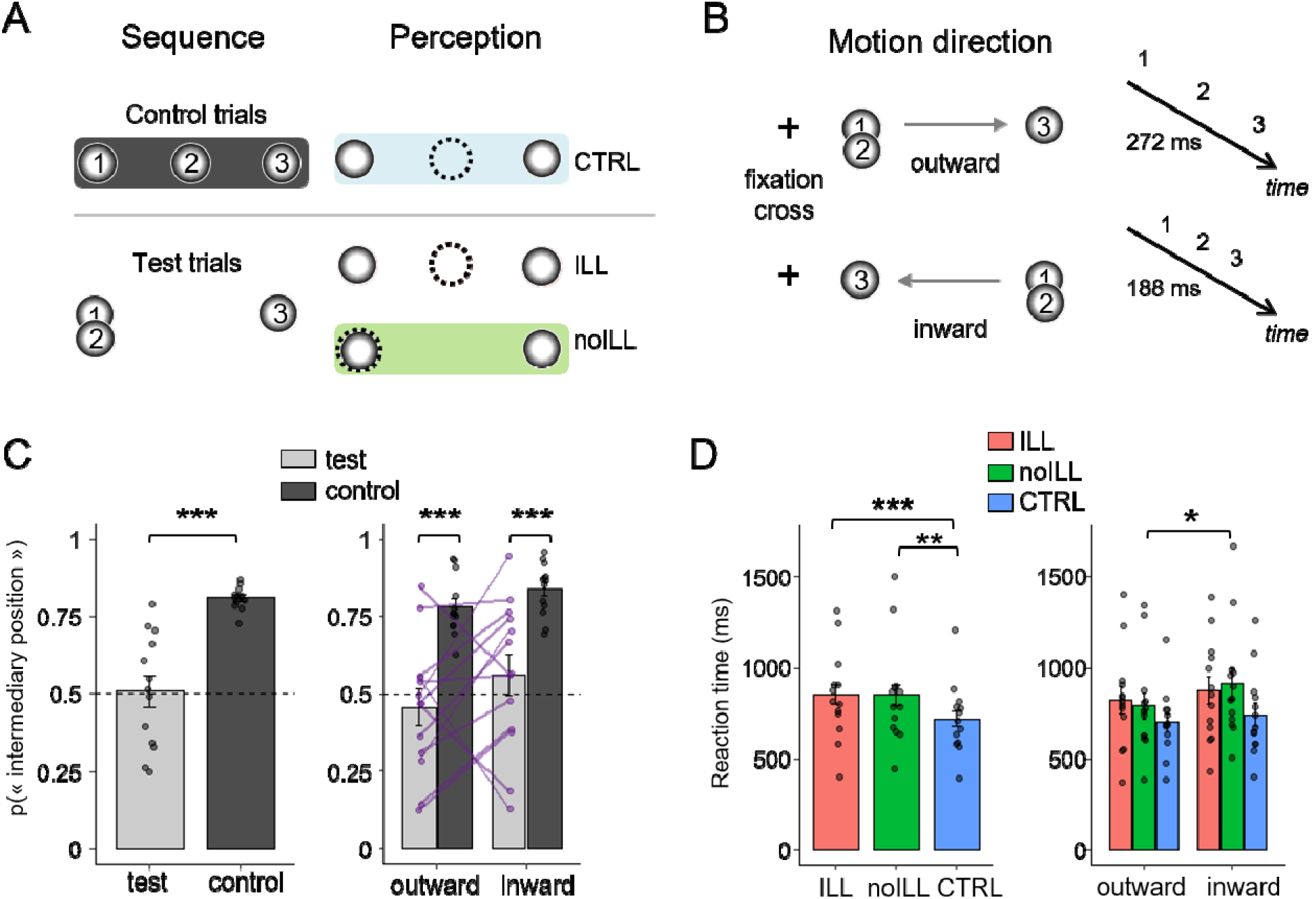
Experimental design and behavioral results. **A.** In test trials, the two first flashes of the sequence were presented at the same spatial location. The second flash was perceived correctly (noILL, in green), or at an intermediary position (ILL, in red). In control trials, the second stimulus was presented and perceived at an intermediary position (CTRL, in blue). Numbers in the circles indicate the temporal order of each flash within the sequence; dashed circle indicates the perceived position of the second flash. **B.** All stimuli were presented on the right horizontal meridian. Two directions for the sequence were possible, to ensure a balanced focus of spatial attention (outward or inward). The SOA was equal to 272ms in outward condition and 188ms in inward condition. **C.** The left panel shows that participants the probability of perceiving the intermediary position in test (light grey) and control trials (dark grey). The right panel shows the same data split following on direction conditions. The purple lines connect individual percentages of illusion in test trials between conditions. The dotted line corresponds to 50% of “intermediary position” response. **D.** In the left panel, reaction times (computed from the onset of the last stimulus) were faster in CTRL trials (blue) than test trials (ILL, red, and noILL, green). The right panel shows that reaction time is faster in outward than inward condition. *** p < .001, ** p< 0.01, * p < .05

The stimulus onset asynchrony (SOA) between the flashes in a sequence was determined through a pilot study (n = 5) so that participants reported 50% of illusory percepts during the presentation of a test sequence. Owing to the fact that attentional strategies can influence the rabbit illusion (Kilgard & Merzenich, 1995), the test condition could follow two possible directions that were centrifugal or centripetal relative to the fixation dot (Fig. 1B). With these two possible motion directions, participants could not predict *where* the next sequence started, ensuring that they payed equal attention to the three spatial locations. The SOA was 188 ms for the centripetal sequence (referred to as “inward” condition) and 272 ms for the centrifugal sequence (“outward” condition). The reasons of this temporal asymmetry remain unclear but it is consistent with the spatial asymmetry observed in the flash-lag effect, another temporal illusion (Kanai, Shet, & Shimojo, 2004).

Participants were told that the locations of the first and the last flashes in the sequence were fixed. They performed a 2-AFC judging whether the second flash was in an intermediate position between the first and last ones. To help them recognize the second flash, it was slightly brighter than the other two flashes: contrast of 1 compared to 0.4 for the first and last flashes. The inter-trial interval was randomly chosen between 800 and 1300 ms following the participants’ response to avoid rhythmical strategy in solving the task.

The experiment was divided into 5 blocks, each containing 60 repetitions of each test condition (inward, outward) and 30 repetitions of the control condition. The order of presentation was randomized within and across individuals. Each test condition was repeated 300 times and each control condition 150 times, leading to 900 trials in total. Each block lasted around 8 min, and participants were encouraged to rest 1-2 min during the breaks.

### Psychophysics analysis

Statistical analyses were conducted with R suite (R core team, 2013). A two-way repeated-measure ANOVA was conducted on the probability of perceiving the second flash in intermediary position with factors of condition (2: test, control) and sequence direction (2: inward, outward). A two-way repeated-measure ANOVA was also conducted on reaction times with factors of perceptual report (3: illusion, no-illusion, control) and sequence direction (2: inward, outward). Reaction times were computed from the onset of the last flash in the sequence. Post-hoc Bonferroni-corrected paired t-tests were computed as well as the Bayes factor for paired t-tests (*BayesFactor* package with R).

### Combined MEG-EEG and anatomical MRI data acquisition

Electromagnetic brain activity was simultaneously with EEG and MEG and recorded in a magnetically shielded room at a 1000 Hz sampling frequency with a high-pass filter of 0.3 Hz. A whole-head Elekta Neuromag Vector View 306 MEG system (Neuromag Elekta LTD, Helsinki) in upright position and equipped with 102 triple sensor elements (one magnetometer and two orthogonal planar gradiometers) and an Easycap EEG cap with 60 sensors were used. Horizontal, vertical electro-oculogram (EOG) and electrocardiogram (ECG) were recorded during the session. The reference electrode for EEG was placed on the top of the participant’s nose. Participants' head position was measured before each block with four head position coils (HPI) placed over the frontal and mastoid areas. The HPI, fiducial points and each EEG electrode were digitalized to help the coregistration with anatomical MRI. A 2min MEG signal without any subject inside the room was recorded the same day. Anatomical landmarks were used to locate the fiducial points (nasion and the left and right pre-auricular areas) and help for the coregistration between electrophysiological and MRI data. After the end of the MEG/EEG session, the participant underwent an anatomical MRI using a 3-T Siemens Trio MRI scanner to provide high-resolution structural brain image. The classical MPRAGE sequence was used as well as the multi-echo FLASH sequence with flip angle at 5° and 30°, which ensure a better separation for skin and skull boundaries during the creation of the head model.

### MEG-EEG data preprocessing

Signal space separation (SSS) was applied to decrease the impact of external noise (Taulu & Kajola, 2005). SSS correction, head movement compensation, and bad channel interpolation were done using MaxFilter Software (Elekta Neuromag) after visual inspection of the raw data. The anatomical brain data were imported with BrainVisa software, segmented with the FreeSurfer image analysis suite (http://surfer.nmr.mgh.harvard.edu/), and then used to generate the 3-layers boundary elements model (BEM) surfaces needed to constrain the forward model.

Structural MRI and MEG/EEG data were realigned during the co-registration step with mrilab (http://mrilab.sourceforge.net) and MNE-Python suite (Gramfort, et al., 2014). Finally, ocular and cardiac artefacts were corrected by rejecting ICA components computed separately for EEG and MEG data, which correlated with ECG/EOG events. Epochs were rejected if their amplitude exceeded a certain threshold (gradiometers: 4000 e−13 T/m, magnetometers: 4 e-12 T, EEG: 250e-6 V). Bad EEG channels were visually detected, and then interpolated.

### MEG-EEG analysis

#### Epoching

The MEG-EEG data analysis was performed with MNE-Python (Gramfort, et al., 2014), Matlab vR2017a (MathWorks) and the Fieldtrip toolbox (Oostenveld, Fries, Maris, & Schoffelen, 2011). Trials whose reaction time was inferior to 200ms, or superior to 4s, from the offset of the second stimulus, were rejected (~2% of rejection rate). The trials were split following the perceptual outcome: in test trials, participants could either perceive the illusion (illusion condition) or perceive the correct stimuli (no-illusion condition); in control trials, only the correctly perceived trials were considered. The data were band-pass filtered between 0.8 and 35 Hz and down sampled to 100 Hz. Epochs were locked on the first stimulus onset, span from −0.3 s to +1.2 s post-stimulus, and no baseline correction was applied. In view of source reconstruction, the number of trials was equalized between responses for each direction. Trials of both directions were then combined: on average, 168 trials (SD = 23) were used when contrasting ILL vs. noILL.

#### Time-frequency analysis

Epochs were defined from −800 to +1 200 ms post-stimulus, downsampled to 333 Hz and low-passed filtered at 160 Hz. Time-frequency analysis was applied on the whole epochs and on each sensor. Power and inter-trial phase coherence (ITC) were calculated using Morlet wavelets for the theta (4-7 Hz, 3.7 cycles) and alpha band (8-12 Hz, 6.7 cycles). As the time-frequency hypotheses focused on alpha pre-stimulus activity, no baseline was applied for the computation of oscillatory activity (Jensen & Mazaheri, 2010; Busch, Dubois, & VanRullen, 2009; Hanslmayr, et al., 2007). Alpha power was computed for combined in- and out-ward directions but also for each direction separately. We restrained the pre-stimulus period from −600 to −200 ms to avoid contamination of post-stimulus responses. Spatiotemporal clustering based on permutation t-test was performed for each sensor and channel type with MNE-Python (cluster threshold = 0.01). Only significant clusters with p-value under .05 were reported. We also correlated the difference in pre-stimulus alpha power between ILL and noILL trials and the individual percentage of illusory perception, for each time point and sensors. The statistical analysis was done with spatiotemporal permutation-based clustering with Fieldtrip functions (cluster threshold = 0.01, minimum neighbors = 2, 4000 repetitions) for each sensor type. Then, we ran a conjunction analysis between directions, which consisted, for each time point and sensor, in keeping the lowest absolute value of correlation coefficient between directions. The conjunction R is then compared to a null distribution computed by shuffling individuals (repetitions = 4000) using spatio-temporal clustering. Alpha power was also computed in source space for each time point with Morlet wavelets (8-12 Hz, 6.7 cycles).

#### Phase analysis

We investigated whether the phase of pre-stimulus alpha activity influenced the perception of a test trial as illusory or veridical, by using the Phase Opposition Sum (POS) index (VanRullen, 2016). This index is based on the inter-trial phase coherences measured over trials of each condition (ITC_ILL_ and ITC_noILL_) and over all trials (ITC_all_): POS = ITC_ILL_ + ITC_noILL_ – 2. ITC_all_.

To investigate the statistical significance of the POS index, we used a permutation procedure (VanRullen, 2016) on the pre-stimulus period (−600 to −200 ms). We first built a null distribution for each individual by shuffling the trial labels and re-computing the associated POS over all sensors and time points (n = 1000). We then built a null distribution at the group level by randomly picking one sample per individual from the previous distributions, averaging the POS data points over individuals and repeating these operations (n = 10 000). A spatio-temporal clustering based on permutation t-test was then performed on the POS index of interest based on ILL and noILL conditions, using the null group distribution. The POS index of interest plotted in Fig. 3A and Fig. 3B was z-scored relative to the group-level null distribution. This analysis was carried out separately for the alpha and theta bands.

#### Source reconstruction

The noise covariance matrix necessary for source reconstruction was estimated from a 5-minute segment of resting state recorded at the beginning of the session. Individual forward solutions were computed using a 3-layers boundary element model constrained by the anatomical MRI. The inverse operator was computed from the noise covariance matrix and the forward model. The dSPM method was used to create the source estimates using a loose orientation constraint (loose = 0.4, depth = 0.8). The estimates were then morphed into the Freesurfer average brain for subsequent group averaging procedure.

To estimates the sources of the correlation analysis in sensors, we correlated the individual difference in alpha power between ILL and noILL trials with the individual percentage of illusory perception. The correlation was done using Spearman’s correlation on each voxel and time point. For visualization, we used the aparc.a2009s parcellation (Destrieux, Fischl, Dale, & Halgren, 2010) and selected the labels containing at least 25 significant voxels at p < 0.01, at any time point included in the significant temporal window found from the sensor analysis. We used a similar procedure for the visualization of POS sources. We computed individual individual alpha POS between ILL and noILL trials, averaged across participants, and selected labels containing at least 25 voxels with a POS value above the 99^th^ percentile value (corresponding to p=0.01) from the null group-level distribution computed in the sensor analysis.

## Results

### Behavioral results

Participants (N = 13) presented with a sequence of 3 flashes were asked to determine the location of the second flash in a 2-AFC task. In test trials, the second flash was presented at the same location than the first flash; in control trials, the second flash was presented at an intermediate spatial location between the first and last flash (Fig. 1A). In both test and control trials, the 3-flash sequences were always presented in the right visual periphery going from left to right (outward) or from right to left (inward, Fig. 1B). In the control condition, participants correctly reported the location of the second flash in 81 % (SD = 1) of the trials (Fig. 1C). In the test condition, the sequences were designed to elicit 50 % of illusory responses and, on average, participants did report an illusory displacement of the second flash in 51 % (SD = 5) of the trials (Fig. 1C). A two-way repeated-measure ANOVA was conducted on the probability of “intermediary position” responses with factors of condition (2: test, control) and direction (2: inward, outward), and subjects as random effect. We found a main effect of experimental condition (F(1,12) = 37.94; p = 5.10^−5^, η^2^_p_ = .76) showing that participants reported the second flash in intermediary position significantly more often in control than in test trials, which was expected by experimental design. We found no significant effects of sequence direction (F(1,12) = 1.83, p = .201, η^2^_p_ = .13) and no interactions between condition and sequence direction (F(1,12) = 1.59, p = .231, η^2^_p_ = .12). Noticeably, the inter-individual variability in illusory perception is large (range = 12-95%) and individual illusory percentages differ between sequence directions (Spearman correlation: R=0.37, p = 0.217, BF = 0.74; see right panel in Fig. 1C).

We also investigated participants’ reaction times, which can provide insights on the chronometry of the spatial reorganization in the *rabbit* illusion. We performed a two-way repeated-measure ANOVA on log-transformed reaction times, with factors of sequence direction (2: inward, outward) and perceptual report (3: illusion, no-illusion, control), and subjects as random effect (Fig. 1D). We found a main effect of perceptual report on reaction times (F(2,12) = 6.750, p = .004, η^2^_p_ = .36) and a marginal effect of sequence direction (F(1,12) = 4.241, p = .062, η^2^_p_ = .26). We found no interactions between both factors (F(2,24) = 1.369, p = .274, η^2^_p_ = .10). A post-hoc Bonferroni-corrected paired t-test showed that reaction times in control trials were on average 130 ms faster than in ILL (p = 10^−4^, BF = 493.0) and 130 ms faster than in noILL (p = .004, BF = 26.7). Slower reaction times in the test as compared to the control trials could be due to an additional perceptual or decisional process having to account with the larger uncertainty of perceptual outputs in the test condition. Although the effect of sequence was marginally significant in the ANOVA analysis, reaction times averaged across all trials were also 71 ms faster in outward than in inward condition (p = .014, BF = 3.1), consistent with previous studies using looming/receding stimuli (Cappe, Thut, Romei, & Murray, 2009; Tyll, et al., 2013).

### Pre-stimulus alpha power correlates with individual tendencies to perceive the illusion

Our working hypothesis was that alpha power could reflect an individual’s slow-speed prior. A strong prior will increase the overall probability to perceive the illusion, so we used the individual percentage of illusory trials as a proxy for this prior. We expected that alpha fluctuations predict whether participants will follow their tendency to perceive the illusion or not. We thus performed a Spearman correlation analysis between the individual likelihood of perceiving the illusion and the alpha power difference between illusion and no-illusion trials. We performed this analysis separately on each direction because the percentage of illusory perception depends on the sequence (Fig. 1C, right panel). A spatiotemporal cluster-based permutation test revealed significant clusters in both sequence conditions located in left central gradiometers (Fig. 2A, inward: −603 to −513 ms, T_sum_ = 234.65, p = 0.029; outward: −585 to −513 ms, T_sum_ = 148.97, p = 0.034). The correlation coefficient for the significant sensors and time points was 0.74 (bootstrap-based CI95% = [0.21, 0.96], BF = 451) and 0.92 (bootstrap-based CI95% = [0.70, 0.99], BF =3855) for the inward and the outward condition, respectively (Fig. 2B). Although the effect appears in similar time points for both conditions, the significant clusters seem spatially distinct between directions. As the spatio-temporal clustering does not specifically address the localization of an effect, we computed the correlation for one direction within the significant cluster of the other direction. The correlations were not significant (inward data in outward cluster: R = −0.43, p = 0.140, bootstrap-based CI95% = [−0.87, 0.14]; outward data in inward cluster: R = 0.07, p = 0.809, bootstrap-based CI95% = [−0.50, 0.59]) Bayes factors indicated anecdotal evidence for a correlation in one condition, and substantial evidence for no correlation in the other one (inward data in outward cluster: BF = 1.739; outward data in inward cluster: BF = 0.229). Additionally, we performed on the whole set of sensors and time points a conjunction analysis based on the assumption of common location, which did not yield any significant effect (p < 0.05). Altogether, these analyses suggested that the correlation arose from distinct sensors for each direction. Finally, we addressed the specificity of the effect by performing the same correlation analysis on theta, beta, low gamma and high gamma frequency bands. No significant cluster (p < 0.05) were found, indicating that the correlation between brain activity and the tendency to perceive the illusion is specific to the alpha band.

**Fig. 2:**
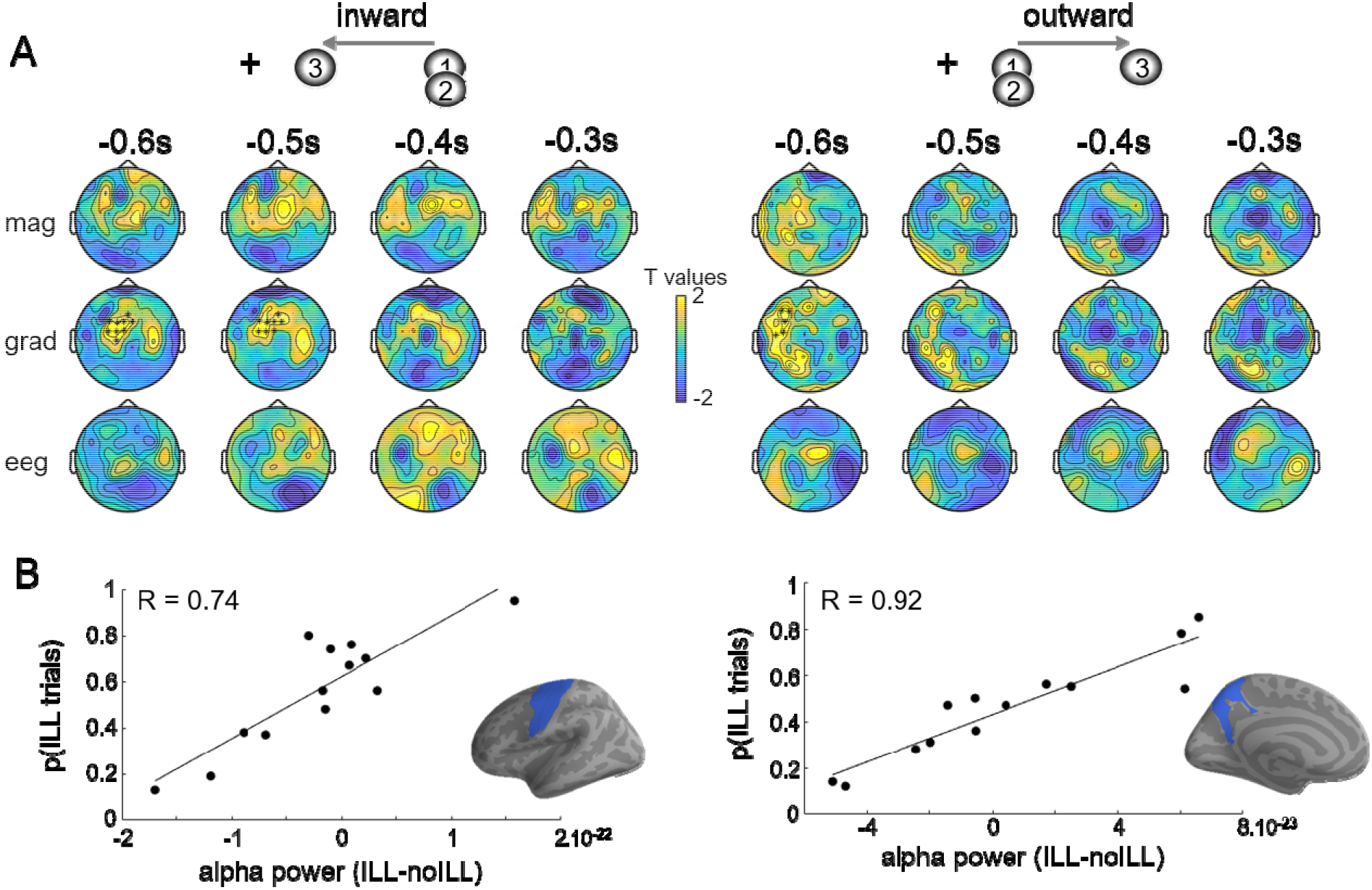
Prestimulus alpha power fluctuations correlate with the individual tendency to perceive the illusion. **A.** Spatio-temporal clustering on the between-individual correlation between the probability of perceiving the illusion and the prestimulus alpha difference ILL-noILL for inward (left) and outward (right) conditions. **B.** The correlation is plotted for significant time points and sensors (bottom pannels). Source reconstruction of the correlation between the probability of perceiving the illusion and the prestimulus alpha difference ILL-noILL for inward (left) and outward (right inset) conditions. Blue labels indicates that at least 25 voxels shows a significant correlation (p < 0.01) within the significant temporal window found in the sensor analysis.

We then used the estimated sources restrained to the significant temporal clusters for each direction and computed the correlation of their amplitude with individual percentages of illusion. The insert in Fig. 2B shows the labels containing at least 25 voxels with a significant correlation at p<0.01. The left precuneus was found for the outward condition (31 significant voxels, R = 0.75 ± 0.04), and the left precentral gyrus (34 significant voxels, R = 0.75 ± 0.04), left central (55 significant voxels, R = 0.75 ± 0.04) and precentral superior sulcus (48 significant voxels, R = 0.75 ± 0.04) were found for the inward condition.

Last, we tested the alternative working hypothesis that prestimulus alpha power directly predicts the perception of the illusion considering that alpha has previously been reported as influencing illusory perception (Keil, Schnitzler, Dijk, Weisz, & Lange, 2014). To test this, we contrasted the power of alpha preceding illusion and no-illusion trials, for combined inward and outward sequences. No significant cluster (p < 0.05) was found after spatio-temporal clustering analysis.

In summary, we found that pre-stimulus alpha power can predict whether an individual will follow or go against his/her tendency to perceive the illusion, not the illusion per se. Additionally, the localization of this effect may depend on the direction of the sequence.

### Pre-stimulus alpha phase predicts the perception of the illusion

Our last hypothesis was that the phase of pre-stimulus oscillations could dissociate illusory trials from non-illusory trials. Based on the literature, we tested two potentially involved frequencies, theta (4-7 Hz) and alpha (8-12 Hz). To address this, we estimated for each frequency the phase consistenc across trials during the pre-stimulus period by using the phase opposition index (POS, see Methods). A significantly positive POS index means that there is a phase concentration at different angles for each tested condition. We ran a spatiotemporal cluster-based test based on a surrogate null distribution to assess the statistical significance of POS between the illusion and no-illusion trials on combined inward and outward sequences (Fig. 3A). We found a significant cluster in the alpha band in gradiometers (−594 to −216 ms, T_sum_ = 20.12, p = 0.043). No significant clusters were found in the theta band (Fig. 3B).

**Fig. 3:**
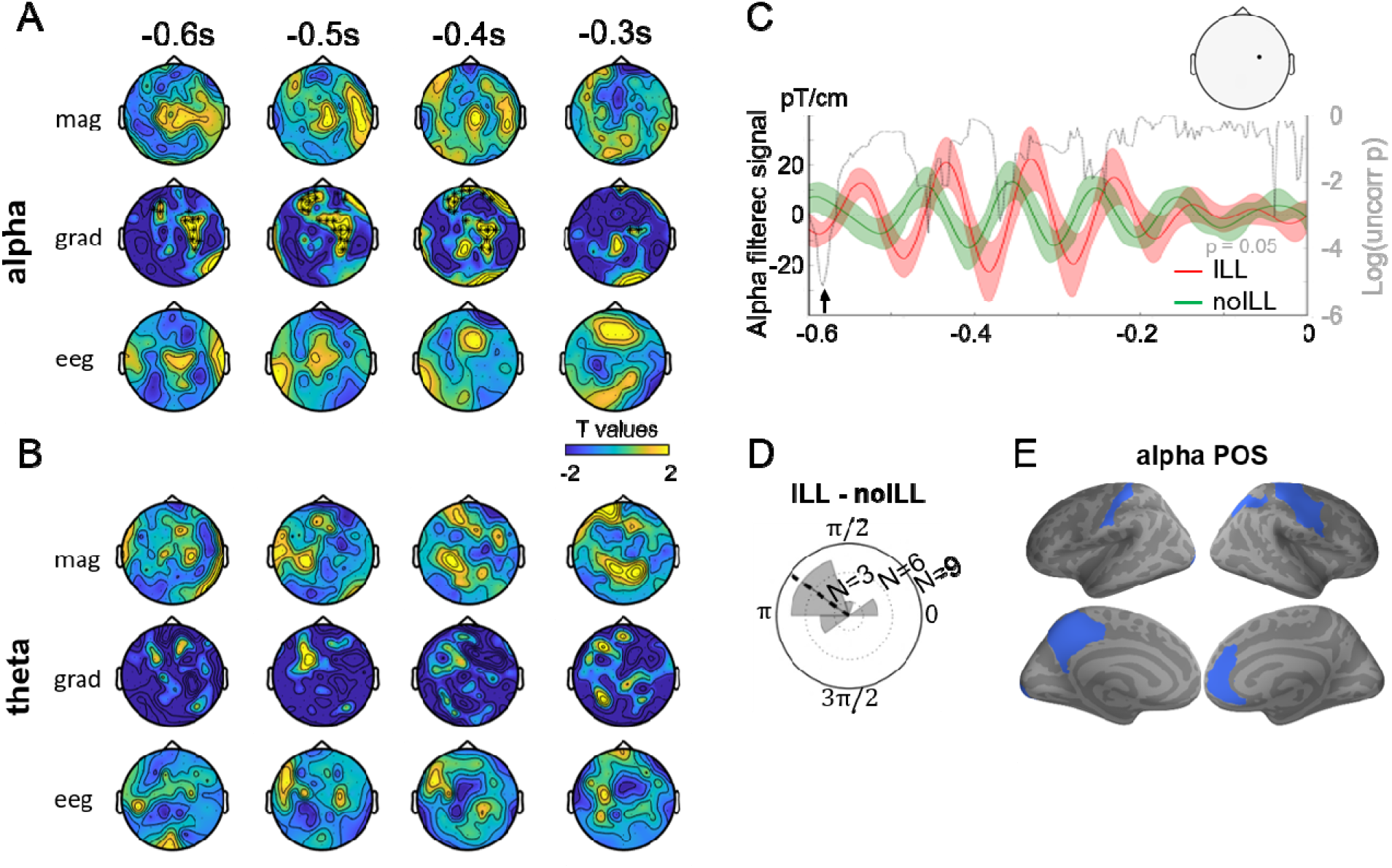
Alpha phase predicts the perception of the illusion. **A.** We investigated whether prestimulus alpha phase predicts the perception of the illusion for combined sequence directions, using POS. A significant cluster was found in the gradiometers. Each topomap show z-scored POS averaged over a 100ms time window. **B.** No significant clusters were found when the theta POS was tested during the pre-sequence period. **C.** For visualization, we selected the channel corresponding to maximal z-scored POS (left topoplot) and filtered the signal in the alpha band, for each response (ILL in red, noILL in green). Shaded areas show SEM. The phase difference between ILL and noILL signals was assessed by a Rayleigh test for each time point. The log of the obtained p-values is plotted in dashed grey line (horizontal dotted line corresponds to p = 0.05). **D.** We selected the time point corresponding to the minimal p-value (showed by the arrow in panel B) and plotted the circular histogram of phase difference between ILL and noILL (mean angle shown by the dashed line). **E.** Source reconstruction of the POS between ILL and noILL trials and the prestimulus alpha difference ILL-noILL, with blue labels indicating that at least 25 voxels have a POS value corresponding to p=0.01 within the significant temporal window found in the sensor analysis.

To visualize this effect, we selected the sensor corresponding to the maximal z-scored POS within the significant cluster, and plotted the grand-average alpha-filtered signal for each response (Fig. 3B). We then computed the instantaneous single-trial phase in the alpha range for this sensor, each response and each participant. We assessed the significance of the phase difference between the illusion and the no-illusion trials using a Rayleigh test on each time point. The uncorrected p-values shown in Fig. 3C (grey curve) reveal a minimal p-value at −582 ms (p = 0.006). The histogram of the phase differences across individuals at that time point showed a mean phase angle of −144° (Fig. 3D), suggesting that perceiving the illusion and not perceiving the illusion can be predicted by opposing alpha phases before the presentation of the visual sequence.

We used source reconstruction and localized the significant cluster in sensors by computing the POS between illusion and no-illusion trials for each voxel and each time point. Fig. 3E shows the labels containing at least 25 voxels with a POS value above a threshold corresponding to p = 0.01 (see Methods). Left precuneus areas (precuneus, 25 voxels; cingulate sulcus, 53 voxels; subparietal sulcus, 28 voxels), left occipital pole (33 voxels) as well as left (postcentral cortex, 29 voxels) and right (postcentral, 34 voxels; precentral, 30 voxels; central cortex, 35 voxels) somatosensory areas, right intraparietal sulcus (28 voxels) and right anterior cingular cortex (43 voxels) were found.

## Discussion

We investigated the role of pre-stimulus oscillatory activity in postdiction herein defined as the mis-localization of an event following the temporal structure of the sequence in which it is embedded (Dennett & Kinsbourne, 1992; Shimojo, 2014). Although alpha power did not directly predict the perception of the illusion, its fluctuations in occipital and parietal areas correlated with the tendency of an individual to perceive the illusion, consistent with the involvement of an internal prior, favoring slow speed motion. We also found that prestimulus alpha phase predicted the perception of the illusion.

### Pre-stimulus alpha activity in occipito-frontal areas relates to prior in postdiction

Based on the assumption that postdiction is driven by prior expectations, and on previous literature indicating that the influence of priors on perception is reflected in low-frequency oscillatory activity (Sherman, Kanai, Seth, & VanRullen, 2016; Mayer, Schwiedrzik, Wibral, Singer, & Melloni, 2016; Samaha, Boutonnet, Postle, & Lupyan, 2018; Nuttida Rungratsameetaweemana & Serences, 2018), we hypothesized that perceptual reports during the rabbit illusion are similarly related to pre-stimulus oscillatory activity. We thus investigated pre-stimulus alpha activity through the prism of a recent line of research which has suggested a specific interpretation of the role of alpha band oscillations during perceptual decisions: alpha may reflect the influence of idiosyncratic biases on perception (Grabot & Kayser, 2019; Grabot, Kösem, Azizi, & Wassenhove, 2017) and may relate to the decision criterion rather than accuracy (Limbach & Corballis, 2016; Iemi, Chaumon, Crouzet, & Busch, 2017; Samaha, Iemi, & Postle, 2017; Craddock, Poliakoff, El-deredy, Klepousniotou, & Lloyd, 2017; Iemi & Busch, 2018). We found that pre-stimulus alpha power fluctuations in parietal and frontal areas reflect the tendency of an individual to perceive the illusion. A participant often reporting the illusory percept has a favorable bias towards the illusion, which will be predicted by high alpha power. On the contrary, a participant often reporting the veridical percept has the opposite bias, predicted by high alpha power. In other words, high alpha power is followed by the perception of the visual sequence in agreement with an individual’s bias, while low alpha power leads to perceive the sequence opposite to one’s own bias. Consistently with recent studies (Limbach & Corballis, 2016; Iemi, Chaumon, Crouzet, & Busch, 2017; Samaha, Iemi, & Postle, 2017), we thus found that alpha power was not related to the physical order of the sequence.

The cerebral sources of this effect were estimated in parietal and frontal areas, consistently with the involvement of top-down signals carried by low-frequency oscillations (Mayer, Schwiedrzik, Wibral, Singer, & Melloni, 2016; Michalareas, et al., 2016). The precuneus, found in the outward condition, is generally related to visuospatial imagery (Cavanna & Trimble, 2006), but also to the maintenance of past stimulus history during multisensory integration (Park & Kayser, 2019). The regions found to be activated in the inward condition, fall within the motor and premotor cortex, known to play a central role in timing perception, especially in the momentarily indexing of ongoing time (Macar, Coull, & Vidal, 2006; Coull, Meck, & Cheng, 2011). Pre-stimulus alpha activity correlating with individual biases was also located in frontal areas, possibly related to cognitive control (Grabot & Kayser, 2019).

The different topographies of the alpha effect between the inward and the outward conditions remain unexplained, as it may be possible that the analysis were underpowered for detecting all the involved cortical regions in both conditions. However, it could also be possible that the relevant motion (slow-speed) priors involved in postdiction differ between movement directions, and recruit different brain regions: looming stimuli were shown to be differently processed compared to receding stimulus in timing and space likely due to their potential ecological importance (Hubbard & Neuhoff, 2018; van Wassenhove, Buonomano, Shimojo, & Shams, 2008).

### Pre-stimulus alpha phase predicts illusory reports

A general hypothesis posits that low-frequency oscillations may segment neural processing into discrete cognitive units (Lakatos, et al., 2005). Alpha phase was found to influence the perception of an upcoming stimulus (Busch, Dubois, & VanRullen, 2009; Mathewson, Gratton, Fabiani, Beck, & Ro, 2009) and was related to cortical excitability (Haegens, Nácher, Luna, Romo, & Jensen, 2011; Kayser, Wilson, Safaai, Sakata, & Panzeri, 2015). These empirical studies have led to the idea that visual information is sampled rhythmically at alpha frequency (VanRullen & Koch, 2003; Samaha & Postle, 2015; Bonnefond & Jensen, 2012; VanRullen, 2016). Importantly, the relationship with alpha phase and perception has usually been studied using a single stimulus. Here, we used a three-flash sequence with a perceptual report concerning the second stimulus and found that the alpha phase preceding the first flash influenced the perceived location of the second flash.

We can speculate about several possible underlying mechanisms and rule out some others. Alpha cycles were proposed to act as integration windows (Cecere, Rees, & Romei, 2015; Milton & Pleydell-Pearce, 2016; Rohe, Ehlis, & Noppeney, 2019; Ronconi, Oosterhof, Bonmassar, & Melcher, 2017), but the flashes used here were separated by ~2-3 alpha cycles (188 or 272 ms), which prevents any stimulus from occurring within the same alpha cycle. An alternative hypothesis could be that the pre-stimulus alpha phase only influences the processing of the first flash, since the first stimulus is likely to phase-reset the ongoing neural oscillations (Romei, Gross, & Thut, 2012; Canavier, 2015). Since alpha phase was found to improve or impair the detection of a visual stimulus (Busch, Dubois, & VanRullen, 2009; Mathewson, Gratton, Fabiani, Beck, & Ro, 2009), a favorable alpha phase could improve the processing of the first stimulus, therefore decreasing the uncertainty on the temporal information of the full sequence. This temporal information would then combine with the internal slow-speed prior, to increase the probability of perceiving the second flash as spatially in-between the first and last ones. More investigation will be needed to determine the specific influence of alpha phase on the sequence processing and perception.

Regarding the localization of this effect, we found implicated regions in occipital cortex, typically observed in studies investigating phase effects on stimulus detection (Mathewson, Gratton, Fabiani, Beck, & Ro, 2009; Sherman, Kanai, Seth, & VanRullen, 2016). The pre-stimulus alpha phase in occipital areas correlated with trial-by-trial variability of the flash-lag effect, another illusion possibly relying on postdictive processes (Chakravarthi & VanRullen, 2012). We noticed that the left precuneus and left premotor areas were implicated in both the alpha power/prior and the alpha phase/accuracy effects, suggesting that they play a key role in postdiction. One fMRI study investigating postdiction also found the implication of premotor and prefrontal cortex, although they used a tactile version of the rabbit *illusion* (Blankenburg, Ruff, Deichmann, Rees, & Driver, 2006). These premotor and prefrontal areas could be involved in general processes underlying postdiction, while the precuneus activity would be specific to visual modality.

Postdictive phenomena have been argued to rely on retro-active processes akin to perceptual reconstruction. Unlike prediction, postdiction is thus considered to take place following the presentation of stimuli. This theoretical argument is at odds with our findings: the initial states of non-linear dynamics (alpha power and phase) can partly determine the perceptual outcome of the rabbit illusion before the presentation of the visual sequence has taken place. Postdictive phenomena may be a special case of predictive mechanisms (Hogendoorn & Burkitt, 2019) in which strong individual priors may shape subjective perception.

